# PARP1 and PARP2 stabilise replication forks at base excision repair intermediates through Fbh1-dependent Rad51 regulation

**DOI:** 10.1101/243071

**Authors:** George E. Ronson, Ann Liza Piberger, Martin R. Higgs, Anna L. Olsen, Grant S. Stewart, Peter J. McHugh, Eva Petermann, Nicholas D. Lakin

## Abstract

PARP1 regulates the repair of DNA single strand breaks (SSBs) generated directly, or during base excision repair (BER). However, the role of PARP2 in these and other repair mechanisms is unknown. Here, we report a requirement for PARP2 in stabilising replication forks that encounter BER intermediates through Fbh1-dependent regulation of Rad51. Whilst PARP2 is dispensable for tolerance of cells to SSBs or homologous recombination dysfunction, it is redundant with PARP1 in BER. Therefore, combined disruption of PARP1 and PARP2 leads to defective BER, resulting in elevated levels of replication associated DNA damage due to an inability to stabilise Rad51 at damaged replication forks and prevent uncontrolled DNA resection. Together, our results demonstrate how PARP1 and PARP2 regulate two independent, but intrinsically linked aspects of DNA base damage tolerance by promoting BER directly, and through stabilising replication forks that encounter BER intermediates.

## INTRODUCTION

The genome of organisms is under constant assault from a variety of agents that cause DNA damage. As such cells have evolved a DNA damage response (DDR) that detects and repairs DNA lesions to restore genome integrity^1^. A central component of this response is signalling DNA damage to effector proteins through post-translational modifications including phosphorylation, ubiquitylation, SUMOylation, acetylation and ADP-ribosylation^2^. This, in turn, regulates a variety of processes such as cell cycle arrest and DNA repair that are critical to maintain genome integrity. The importance of these pathways is underscored by the observations that defects in these pathways leads to chromosome instability and a variety of pathologies, including increased cancer risk.

ADP-ribosyltransferases (ARTDs), or poly(ADP-ribose) polymerases (PARPs), attach ADP-ribose onto target proteins either as single units or polymer chains by mono-ADP ribosylation (MARylation) or poly-ADP ribosylation (PARylation) respectively^3^. Of the 17 human genes containing predicted ARTD catalytic domains^4^, several have been identified as primary sensors of DNA damage^5^. PARP1, the founding member of the ARTD family, senses DNA single strand breaks (SSBs) induced either directly, or as a consequence of processing DNA lesions during the base excision repair (BER) pathway^6^. PARP1 becomes activated upon binding SSBs and PARylates a variety of substrates to promote the accumulation of XRCC1 at damage sites that subsequently acts as a scaffold to assemble repair factors at the break^7, 8, 9, 10^. PARP1 also regulates pathways other than SSB repair (SSBR) including replication fork progression and re-start^11, 12, 13^, although the mechanisms of this regulation are unclear. It also promotes alternative non-homologous end-joining (alt-NHEJ), a pathway activated in the absence of core NHEJ^14, 15^. Whilst PARP1 has also been implicated in canonical NHEJ^13, 16^, PARP3 promotes this pathway by facilitating accumulation of repair factors such as APLF and Ku at damage sites^17, 18, 19^.

Although PARPs regulate several different DNA repair mechanisms, it is unclear how overlapping functions between these enzymes promotes cell viability in the face of genotoxic stress. For example, PARP2 has been implicated in repair of DNA base damage^20, 21^ and redundancy between PARP1 and PARP2 is implied by embryonic lethality of *parp1^-/-^parp2^-/-^* mice^20^. However, the role of PARP2 in regulating DNA repair and its relationship to PARP1 in this process remain unknown. Moreover, whether disruption of PARP-dependent SSBR results in elevated levels of DNA damage that are channelled through alternate repair mechanisms, how these lesions are processed, and whether this is also regulated by PARPs is unclear. Given PARP1 and PARP2 are both targets for inhibitors being used to treat tumours with defects in homologous recombination (HR)^22, 23^, unravelling these complexities will be important not only for understanding the mechanistic basis of DNA repair, but also refining the use of PARP inhibitors (PARPi) in the clinic and guiding the development of novel PARPi with new mechanisms of action.

Here we address these questions by disrupting PARP1 and PARP2 alone or in combinations and assessing the impact of these manipulations on the repair of DNA base damage. We identify that PARP1 and PARP2 are redundant in BER and allow cells to tolerate DNA base damage induced by methyl methanesulfonate (MMS). Surprisingly, we find that redundancy between PARP1 and PARP2 does not extend to synthetic lethality with HR-deficiency and that loss of PARP1 is the major driver of this phenotype. Moreover, in the absence of PARP1, PARP2 is required for optimal resolution of MMS-induced DNA damage during DNA replication, independent of its role in BER, through stabilising HR proteins at sites of replication stress to protect stalled and/or damaged forks against uncontrolled nucleolytic resection.

## RESULTS

### PARP1 and PARP2 are redundant in base excision repair

In order to understand the contributions of different PARP family members in regulating DNA repair, we generated a number of cell lines deficient for PARPs in U2OS cells, starting with PARP1 (Fig. 1a and Supplementary Fig. 1). Consistent with the role of PARP1 in BER and SSBR^6^, *parp1∆* cells exhibit sensitivity to the DNA alkylating agent MMS and the oxidative DNA damage agent H_2_O_2_ (Fig. 1a and Supplementary Fig. 2a). However, although MMS-induced nuclear ADP-ribosylation is significantly reduced in *parp1∆* cells, it is not completely defective (Fig. 1d), and the PARP inhibitor olaparib further sensitises *parp1∆* cells to this agent (Fig. 1b). Taken together, these data indicate that whilst PARP1 is required for the cellular response to DNA base alkylation, in its absence an additional PARP(s) responds to this variety of DNA damage.

**Figure 1:**
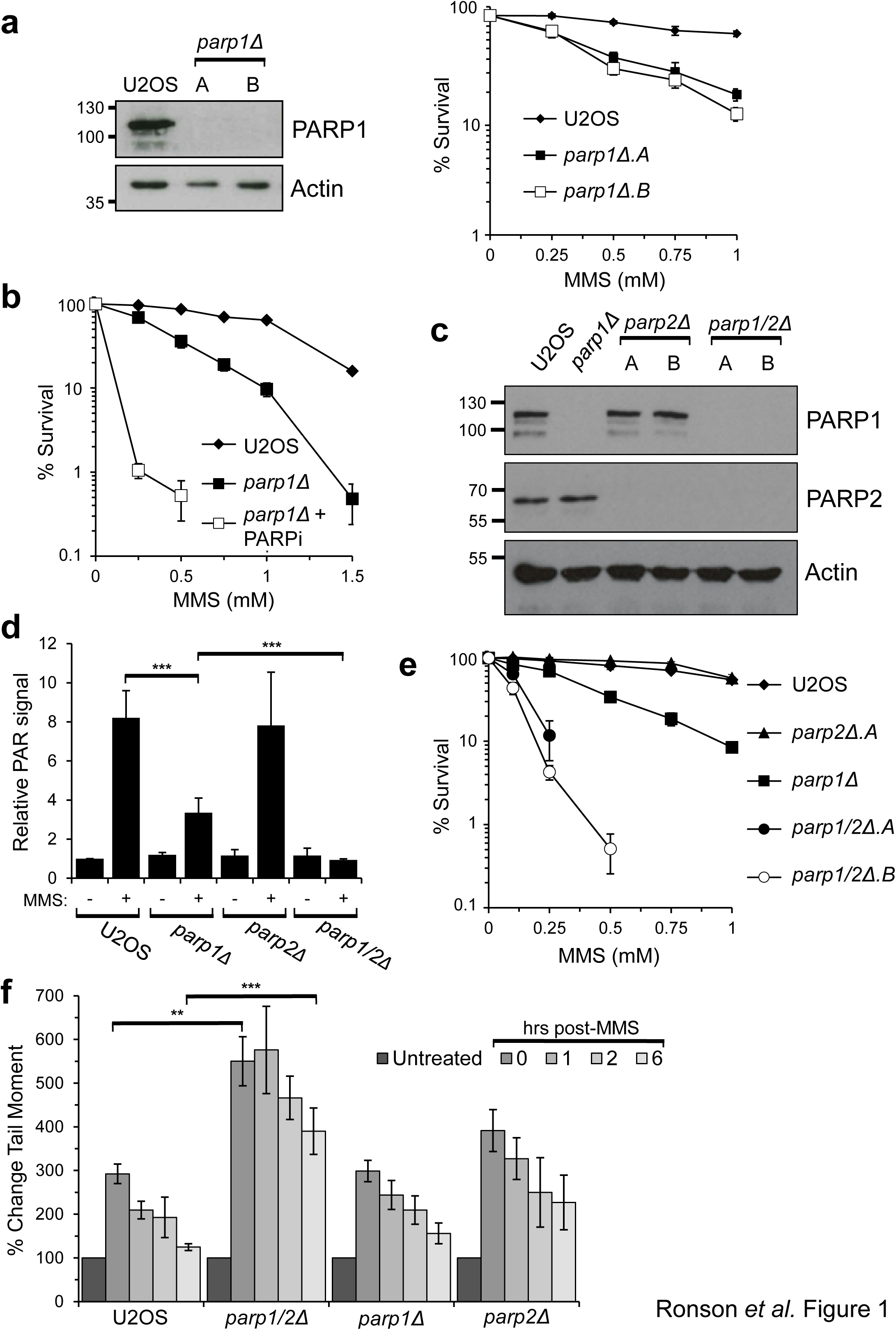
PARP1 and PARP2 are redundant in the cellular response to MMS-induced DNA damage. **a.** Whole cell extracts were prepared from U2OS or two independent *parp1∆* cell lines and Western blotting performed with the indicated antibodies (left panel). U2OS or *parp1∆* cell lines were exposed to MMS and cell survival assessed by clonogenic assays (right panel). Error bars represent the standard error of the mean (SEM) from three independent experiments. **b.** U2OS or *parp1∆* cell lines, with or without exposure to Olaparib (PARPi), were exposed to MMS and cell survival assessed by clonogenic assays. Error bars represent the SEM from three independent experiments. **c.** Whole cell extracts were prepared from U2OS, *parp1∆, parp2∆* and *parp1/2∆* cells and Western blotting performed using the indicated antibodies. **d.** The indicated cell lines were left untreated (-) or exposed to 1.5 mM MMS (+) for 1 hour and nuclear ADP-ribosylation analysed by immunofluorescence. Error bars represent the SEM from three independent experiments. **e.** U2OS, *parp1∆, parp2∆ or parp1/2∆* cell lines were exposed to MMS and cell survival assessed by clonogenic assays. Error bars represent the SEM from three independent experiments. **f.** Cells were treated with 250 μM MMS for 1 hour, before recovery in fresh media. Samples were taken at the indicated times post-treatment and the alkaline comet assay used to reveal strand breaks and alkali-labile sites. Comet data was normalised to the untreated sample. Error bars represent mean values ± SD from at least six independent experiments.

Given olaparib is also able to inhibit PARP2^22, 23^, we assessed whether this ARTD also contributes towards the cellular response to MMS by disrupting the *PARP2* gene either alone or in combination with *PARP1* to generate *parp2∆* and *parp1/2∆* cell lines (Fig. 1c and Supplementary Fig. 3). Disruption of *PARP2* alone has little impact on MMS-induced nuclear ADP-ribosylation or the sensitivity of cells to DNA damage induced by this agent (Fig. 1d,e). Strikingly, the residual ADP-ribosylation observed in *parp1∆* cells is lost upon deletion of PARP2 (Fig. 1d) and *parp1/2∆* cells are more sensitive to MMS than when *PARP1* is disrupted alone (Fig. 1e). Importantly, depletion of PARP2 by siRNA also sensitises *parp1∆* U2OS and *parp1∆* RPE-1 cells to MMS (Supplementary Fig. 4 and 5), illustrating a reproducible phenotype using two independent gene depletion strategies in different cell lines. To establish whether redundancy between PARP1 and PARP2 in tolerance of cells to MMS is reflected in their ability to repair SSBs generated during BER, we performed alkaline comet assays following exposure of *parp1∆, parp2∆* and *parp1/2∆* cells to MMS to directly assess the kinetics of SSB resolution (Fig. 1f). Disruption of *PARP1* or *PARP2* alone has no dramatic impact on repair of SSBs generated in response to MMS. In contrast, *parp1/2∆* cells show a significant increase in the induction of DNA strand breaks in response to MMS that persist up to 6 hours after removal of the genotoxin. Taken together, these data indicate that in the absence of PARP1, PARP2 plays a key functional role in signalling DNA damage to promote SSB resolution and cell survival in response to MMS.

### PARP1, but not PARP2 or PARP3 is synthetic lethal with HR

Synthetic lethality between PARP inhibition and HR-deficiency is mediated through targeting PARPs at DNA SSBs^24, 25^. Given our findings that PARP1 and PARP2 are redundant in BER, we considered whether this relationship extends to synthetic lethality with HR-deficiency. Currently available PARPi target PARP1, PARP2 and PARP3^22, 23^. Therefore, we also wished to establish which combination of DNA damage responsive PARP disruptions exhibits the strongest impact on cell viability in combination with HR dysfunction. To address these questions, we inhibited HR in different PARP-deficient backgrounds using B02, a small molecule inhibitor of Rad51 that disrupts its binding to single stranded DNA during nucleofilament formation *in vitro* and *in vivo*, resulting in defective HR^26^. As predicted, B02 inhibits Rad51 foci formation in U2OS cells exposed to MMS or the DSB-inducing agent phleomycin (Supplementary Fig. 6). Consistent with conservation of the synthetic lethal relationship between HR-deficiency induced by B02 and PARP inhibition, cells treated with PARPi are more sensitive to B02 than untreated cells (Fig. 2a). Additionally, PARPi sensitivity is suppressed by the addition of DNA-PKcs inhibitors (DNA-PKi; Fig. 2a), an observation that was originally made in *BRCA2^-/-^* models^27^, further validating this as an appropriate assay to assess the impact of different PARP gene disruptions on cell viability in an HR-deficient background.

**Figure 2:**
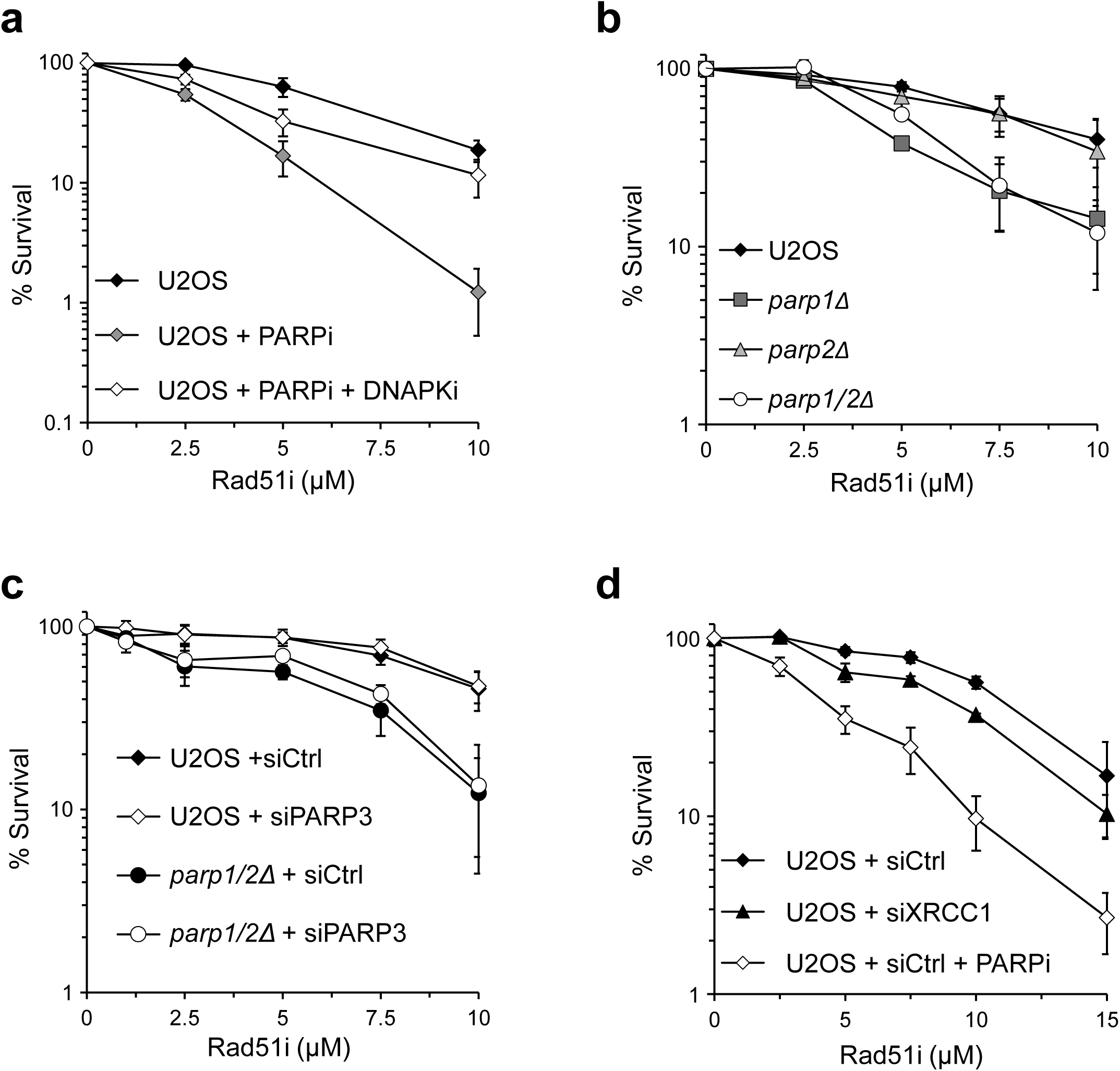
Loss of PARP1, but not PARP2 or PARP3, is synthetic lethal with HR inhibition. **a.** U2OS cells were exposed to B02, with or without Olaparib (PARPi) or Nu7441 (DNA-PKi), and cell viability assessed by clonogenic survival assay. Error bars represent the SEM from three independent experiments. **b.** U2OS, *parp1∆, parp2∆ orparp1/2∆* cell lines were exposed to B02 and cell viability assessed by clonogenic survival assay. Error bars represent the SEM from three independent experiments. **c and d.** U2OS and *parp1/2∆* cell lines, with or without siRNA depletion of PARP3 (C), or U2OS cells with or without XRCC1 siRNA depletion (D), were exposed to B02 and cell viability assessed by clonogenic survival assay. Error bars represent the SEM from three independent experiments.

Whilst *parp1∆* cells are more sensitive than U2OS cells to HR inhibition, *parp2∆* cells showed no sensitivity to B02. In striking contrast to the relationship observed with MMS, *parp1/2∆* cells are no more sensitive to B02 than *parp1∆* cells (Fig. 2b). The absence of PARP3 had no impact on the survival of cells following exposure to B02 irrespective of the presence or absence of PARP1 and PARP2 (Fig. 2c). Additionally, U2OS and *parp3∆* cells can be equally sensitised to B02 by treatment with olaparib (Supplementary Fig. 7d), further confirming that PARP3 disruption is not toxic in combination with HR-deficiency. The lack of redundancy between PARP1 and PARP2 in cell viability with HR-inhibition is in striking contrast to the context of SSB resolution during BER (Fig. 1e,f), suggesting that SSBR defects *per se* are not toxic with HR dysfunction. In support of this hypothesis, depletion of the key BER/SSBR protein XRCC1 is unable to dramatically sensitise cells to B02 (Fig. 2d). Taken together, these data indicate that whilst PARP1 and PARP2 are redundant in terms of BER/SSBR, PARP1 is the major PARP whose disruption is toxic with HR deficiency.

### PARP1 and PARP2 promote repair of DNA damage during S-phase

Whilst *parp1∆* cells are sensitive to MMS, resolution of breaks generated during BER is relatively normal in these cells (Fig. 1f). Additionally, although PARP2 promotes tolerance of cells to MMS in the absence of PARP1, this redundancy is not evident following exposure of cells to H_2_O_2_, an agent that induces DNA SSBs directly (Supplementary Fig. 2b). Together, these data suggest the redundancy between PARP1 and PARP2 in the cellular response to MMS extends beyond a simple role of these enzymes in regulating BER. BER is initiated by recognition and excision of the damaged base using DNA glycosylases. The resulting apurinic (AP) site is subsequently cleaved by AP-endonuclease 1 (APE1), leading to a DNA strand break that is channelled through XRCC1 and PARP1-dependent SSBR^6^. Although BER is a major mechanism for repair of alkylated base damage^6, 28^, MMS also induces DNA damage during S-phase when SSBs created by APE1 cleavage are converted to DSBs by active replication forks^29, 30^. Therefore, we also considered the role of PARP1 and PARP2 in resolution of S-phase associated DNA damage induced by MMS.

To address this question we labelled cells undergoing S-phase with the base analogue 5-ethynyl-2'-deoxyuridine (EdU) to monitor DNA damage either in G1/G2 cells, or cells undergoing DNA replication at the time of DNA damage induction (Fig. 3a). Consistent with a previous report^30^, we observe an increase in γ-H2AX foci in cells undergoing DNA replication at the time of MMS exposure, with no detectable induction of γ-H2AX foci in G1 or G2 cells at the doses employed in this assay (Fig. 3b). No significant increase in MMS-induced γ-H2AX foci is apparent in replicating *parp1∆* or *parp2∆* cells compared to U2OS cells. In contrast, the induction of γ-H2AX foci is significantly elevated in *parp1/2∆* cells, indicating an increase in the levels of S-phase associated DNA damage. To understand whether loss of PARP1 and/or PARP2 affects repair of this damage, we assessed the kinetics of γ-H2AX foci decay at times following removal of MMS. Twelve hours after induction of DNA damage U2OS cells show a decay in the number of γ-H2AX foci, with levels approaching those observed in untreated cells (Fig. 3c). Whilst decay of γ-H2AX foci in *parp2∆* cells is comparable to parental U2OS, *parp1∆* cells show a mild delay in the repair of this damage. Strikingly however, *parp1/2∆* cells exhibit a significant persistence of γ-H2AX foci above that observed in *parp1∆* cells, indicating resolution of DNA damage is compromised (Fig. 3c).

**Figure 3:**
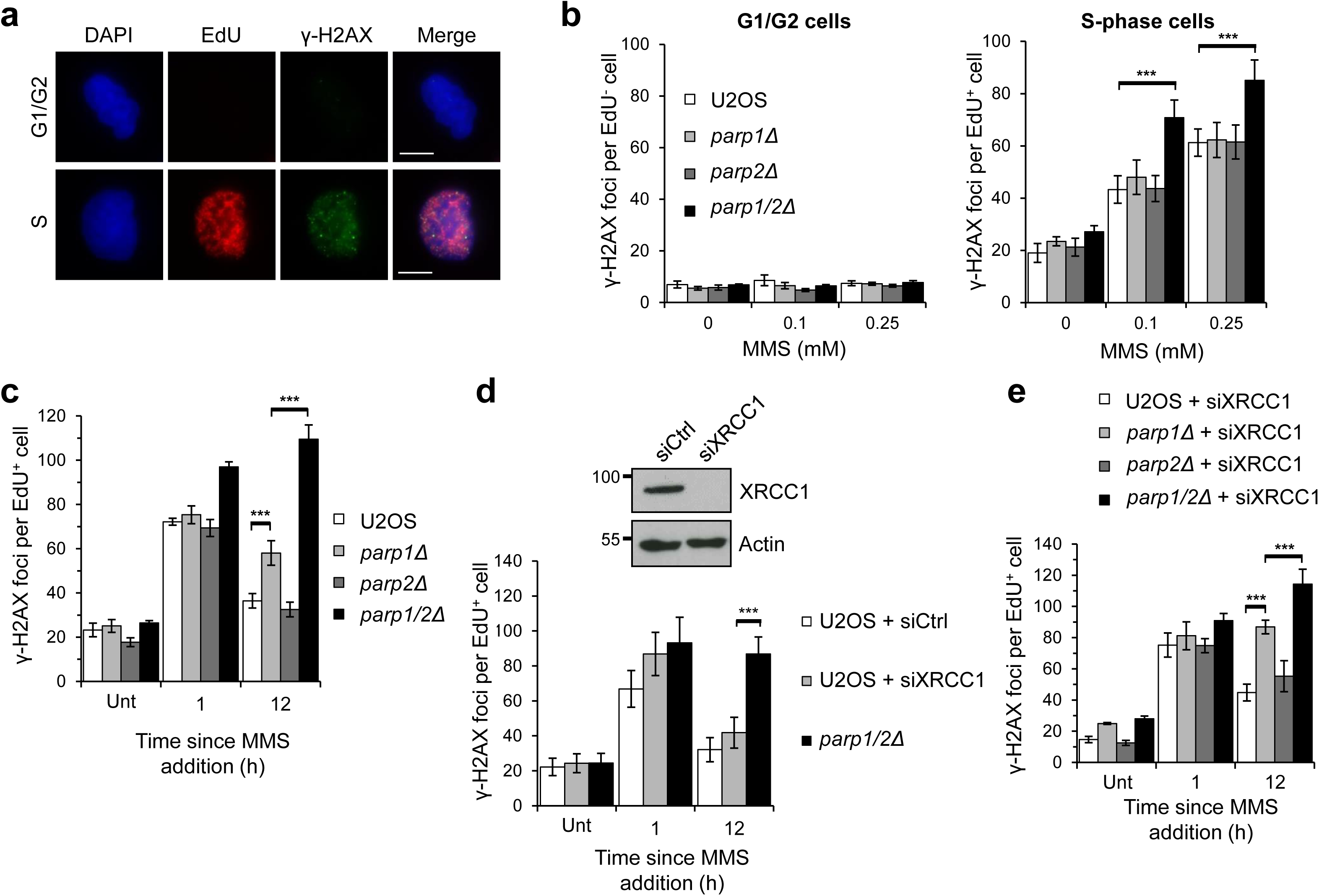
PARP1 and PARP2 function in the resolution of replication-associated damage independently of BER. **a.** U2OS cells were treated with 1 μM EdU and 0.25 mM MMS for 1 hour, before pre-extraction and fixation. EdU and γ-H2AX were detected by immunofluorescence. Representative images of EdU-positive and EdU-negative nuclei are shown. Scale bars represent 10 μm. **b.** The indicated cell lines were treated as in (A) and following immunofluorescence γ-H2AX foci quantified in G1/G2 phase (EdU-negative) and S-phase (EdU-positive) cells. Error bars represent the SEM from four independent experiments. **c.** The indicated cell lines were treated with 1 μM EdU (Unt), or with 1 μM EdU and 0.25 mM MMS for 1 hour before recovery in fresh media, and γ-H2AX foci quantified by immunofluorescence in EdU-positive cells. Times indicated show hours since addition of EdU and MMS. Error bars represent the SEM from four independent experiments. **d.** U2OS cells transfected with control or XRCC1 siRNA, or *parp1/2∆* cells were treated as in (C) and γ-H2AX foci quantified in EdU-positive cells (lower panel). Knockdown of XRCC1 in U2OS cells was confirmed by Western blotting using the indicated antibodies (upper panel). Times indicated show hours since addition of EdU and MMS. Error bars represent the SEM from four independent experiments. **e.** The indicated cell lines transfected with XRCC1 siRNA were treated as in (C) and γ-H2AX foci quantified in EdU-positive cells. Times indicated show hours since addition of EdU and MMS. Error bars represent the SEM from three independent experiments.

Given that disruption of PARP1 and PARP2 results in delayed ligation of DNA breaks (Fig. 1f), a potential explanation for the persistent γ-H2AX foci in replicating *parp1/2∆* cells is that the greater number of unrepaired SSBs in these cells results in elevated levels of S-phase associated DNA damage that requires a longer time for repair. Alternatively, PARP1 and/or PARP2 may play a BER-independent role in resolving DNA damage induced by MMS specifically during DNA replication. To distinguish between these two possibilities we depleted XRCC1, a protein whose knock-down or mutation delays BER and sensitises cells to MMS^31, 32, 33, 34^, and assessed whether this could recapitulate the delayed γ-H2AX decay in a PARP-independent manner. The levels of γ-H2AX foci induced in replicating cells following exposure to MMS increases slightly in XRCC1-depleted cells, presumably through active replication forks encountering unrepaired SSBs (Fig. 3d). Importantly, decay of these foci is relatively normal, resulting in significantly fewer foci at later time points compared to *parp1/2∆* cells, indicating a BER defect alone cannot explain the persistent γ-H2AX foci that result from *PARP1* and *PARP2* gene disruption. To establish whether PARP1, PARP2 or both are required for this BER-independent replication repair event, we depleted XRCC1 in U2OS, *parp1∆, parp2∆*, and *parp1/2∆* cells and assessed γ-H2AX decay after MMS treatment (Fig. 3e). XRCC1-depleted *parp1∆* cells show a marked delay in the resolution of γ-H2AX. Decay of γ-H2AX foci in XRCC1-depleted *parp2∆* cells are comparable to U2OS cells. However, depletion of XRCC1 in *parp1/2∆* cells compromises the decay of γ-H2AX above that observed in XRCC1-depleted *parp1∆* cells. Taken together, these data indicate a role for PARP1 in facilitating replication-associated repair independently of BER and that in its absence PARP2 can contribute to resolution of MMS-induced DNA damage during S-phase.

### PARP1 and PARP2 regulate Rad51 assembly at replication forks

Given the importance of PARP1 and PARP2 in resolving DNA damage during S-phase, we next assessed whether MMS stalls active replication forks and the impact of disrupting PARP1 and PARP2 on their restart. To achieve this, we used DNA fibre analysis in conjunction with a labelling protocol that allowed us to monitor stalled replication tracts induced by MMS and their subsequent recovery following its removal. Ongoing forks were labelled with CldU before stalling by addition of MMS. Following washout of MMS and CldU, cells were released into fresh IdU-spiked media, thereby ensuring that only ongoing and restarted replication forks incorporate the second label whilst the remaining stalled forks appear as CldU-only tracts (Fig. 4a). MMS induced replication fork stalling, with the number of stalled forks decreasing following the removal of MMS, indicating replication fork recovery (Fig. 4a). Interestingly, whilst a comparable number of stalled forks are apparent in *parp1∆* and *parp2∆* cells relative to U2OS cells, *parp1/2∆* cells showed a significantly lower amount of fork stalling at the earliest time point of release (Fig. 4a). Although the number of stalled forks were lower in *parp1/2∆* cells throughout the recovery period, the kinetics of restart were similar to U2OS, *parp1∆* and *parp2∆* cells. Taken together, these data indicate that loss of PARP1 and PARP2 impair fork stalling at MMS-damaged replication templates rather than restart of stalled and/or damaged replication forks.

**Figure 4:**
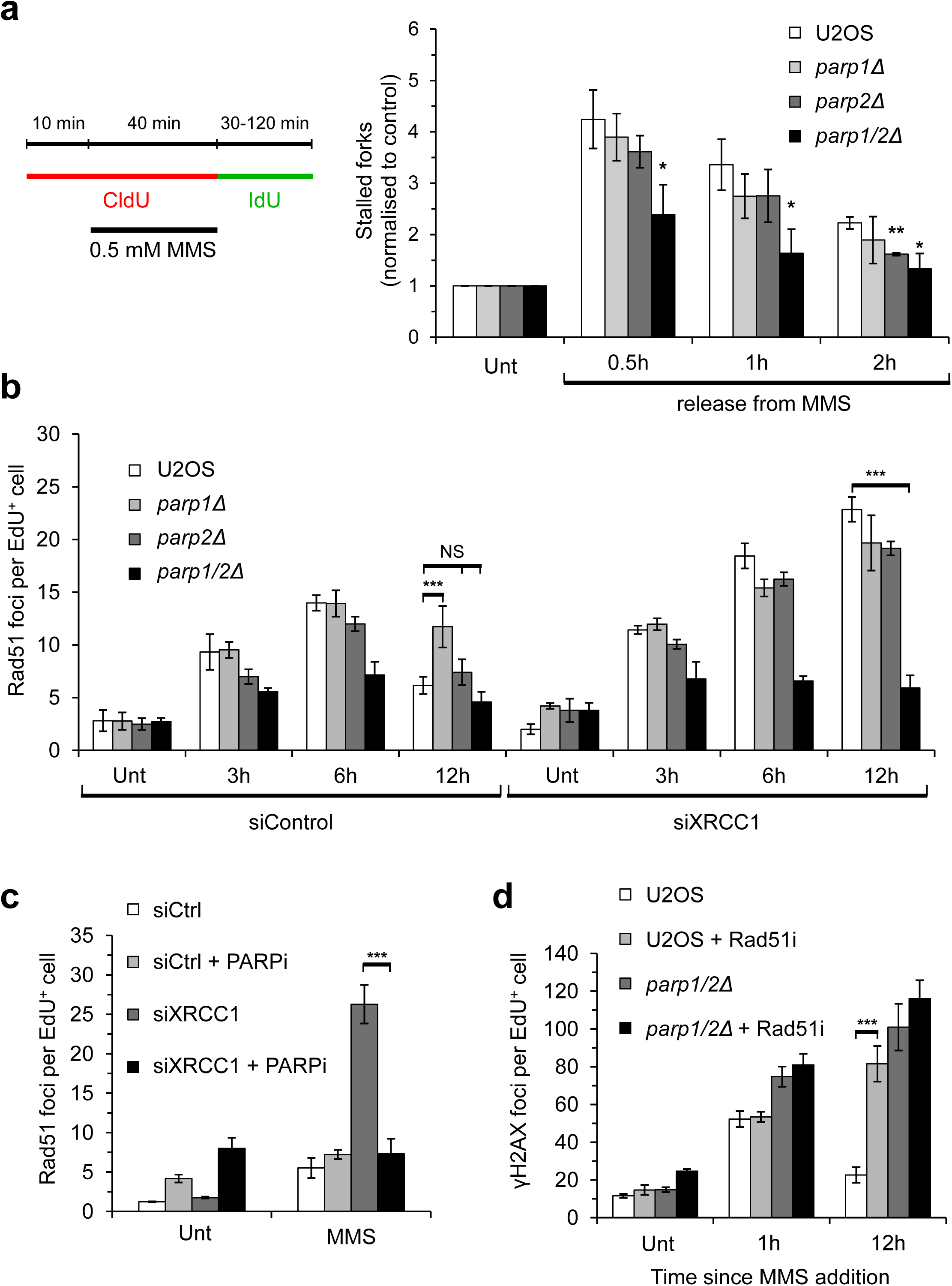
Assembly of Rad51 at damaged replication forks is compromised in the absence of PARP1 and PARP2. **a.** DNA fibre analysis was performed in the indicated cells after MMS exposure and recovery for the times shown. The number of stalled forks (red-only tracts) was determined as a ratio of all red-labelled replication structures. These values were subsequently normalised to the equivalent ratio in the corresponding non-treated sample. Error bars represent the SEM from at least three independent experiments. Statistically significant difference from the normalised ratio of stalled forks in parental U2OS cells at the respective time points is indicated. **b.** The indicated cell lines transfected with control or XRCC1 siRNA were treated with 1 μM EdU (Unt), or with 1 μM EdU and 0.5 mM MMS for 1 hour before recovery in fresh media, and Rad51 foci quantified in EdU-positive cells. Times indicated show hours since addition of EdU and MMS. Error bars represent the SEM from three independent experiments. **c.** U2OS cells transfected with control or XRCC1 siRNA were treated as in (B) and Rad51 nuclear foci analysed in EdU positive cells. Cells were fixed 12 hours after addition of EdU and MMS. Error bars represent the SEM from three independent experiments. **d.** U2OS or *parp1/2∆* cells, in the presence or absence of 50 μM B02 (Rad51 i), were treated as in (3C) and γ-H2AX foci quantified in EdU-positive cells. Times indicated show hours since addition of EdU and MMS. Error bars represent the SEM from three independent experiments.

An inability to slow/stall replication forks in response to a variety of DNA damaging agents is a characteristic outcome of disrupting genes involved in HR^12, 35, 36^. Given disruption of PARP1 and PARP2 results in a similar reduced ability to stall replication forks in response to MMS (Fig. 4a), a phenotype also observed in Rad51-depleted U2OS cells (Supplementary Fig. 8), we next considered whether this effect is a consequence of defective HR. A central component of the HR pathway is assembly of Rad51 at resected DNA termini to stabilise stalled/collapsed replication forks to facilitate their restart once repair is complete^37^. Therefore, we assessed the formation and decay of Rad51 foci in our *parp∆* cell lines at time points following a transient exposure to MMS. Parental U2OS, *parp1∆* and *parp2∆* cells exhibit robust induction of Rad51 foci in EdU positive cells following a transient exposure to MMS, peaking at 6 hours (Fig. 4b). However, whilst decay of these foci over time is similar between U2OS and *parp2∆* cells, they persist in *parp1∆* cells indicating delayed kinetics of DNA repair and/or that elevated levels of DNA damage are channelled through a Rad51-dependent repair mechanism. Interestingly, although we observed a slight induction and decay of Rad51 foci in *parp1/2∆* cells, this was at a lower level to what we observed in the absence of PARP1 or PARP2 (Fig. 4b), suggesting these cells have difficulties in assembly and/or maintenance of Rad51 at damaged replication forks induced by MMS.

To better understand this observation, we also considered the role of PARP1 and PARP2 in regulating HR by assessing Rad51 foci in our *parp∆* cell lines when SSBR has been compromised independently of PARP status. Depletion of XRCC1 results in elevated levels of Rad51 foci that continue to accumulate up to 12 hours following removal of MMS in U2OS, *parp1∆* and *parp2∆* cells (Fig. 4b). This observation was also recapitulated in replicating RPE-1 and MRC5 cells depleted of XRCC1 following exposure to MMS (Supplementary Fig. 9), indicating that disruption of SSBR/BER leads to elevated levels of S-phase associated DNA damage that is channelled through Rad51-dependent repair in several different cell lines. Strikingly, this increase in Rad51 foci is compromised in *parp1/2∆* cells, or U2OS cells treated with olaparib (Fig. 4b,c). PARPi similarly decreases the number of strong MMS-induced Rad51 foci in MRC5 and RPE-1 cells (Supplementary Fig. 9), as does depletion of PARP2 in *parp1∆* RPE-1 cells (Supplementary Fig. 10). Together, these data indicate novel redundancy between PARP1 and PARP2 in the assembly and/or stabilisation of Rad51 at sites of stalled/damaged replications forks in multiple cell lines. Moreover, whilst inhibition of Rad51 using B02 induces a striking increase in persistent γ-H2AX foci following a transient exposure of U2OS cells to MMS, it does not induce a similarly large increase in *parp1/2∆* cells (Fig. 4d). These data suggest that Rad51 and PARP1/PARP2 function in the same pathway with regards to the resolution of S-phase associated DNA damage induced by MMS, providing an explanation for the persistent γ-H2AX foci observed in *parp1/2∆* cells.

### PARP1 and PARP2 stabilise Rad51 at damaged replication forks

PARP1 has previously been implicated in the restart of stalled replication forks through a mechanism that is dependent on Mre11, suggesting that DNA end resection is a critical control point regulated by PARPs^11, 13^. To investigate this possibility in the context of replication stress induced by DNA base alkylation, we assessed ssDNA formation by monitoring RPA foci in EdU-positive cells following a transient exposure to MMS. Similar to Rad51 foci (Fig. 4b), U2OS and *parp2∆* cells exhibit a transient induction of RPA foci following exposure to MMS, whilst they persist in *parp1∆* cells (Fig. 5a). Depletion of XRCC1 induces a continued accumulation of RPA foci in U2OS, *parp1∆* and *parp2∆* cells (Fig. 5b), further supporting the notion that defective SSBR/BER increases DNA damage that is channelled through HR. Strikingly, RPA foci and RPA phosphorylation on serine-4/8 (S4/8) are increased in *parp1/2∆* cells above that observed upon depletion of XRCC1 alone, or in combination with *parp1* and *parp2* gene disruption (Fig. 5a-c), indicating increased levels of ssDNA in these cells above that attributable to defects in SSB/BER. Taken together, these data indicate that although PARP1 and PARP2 are redundant in the assembly of Rad51 at sites of stalled/damaged replication forks induced by MMS, this is not regulated through controlling DNA end-resection.

**Figure 5:**
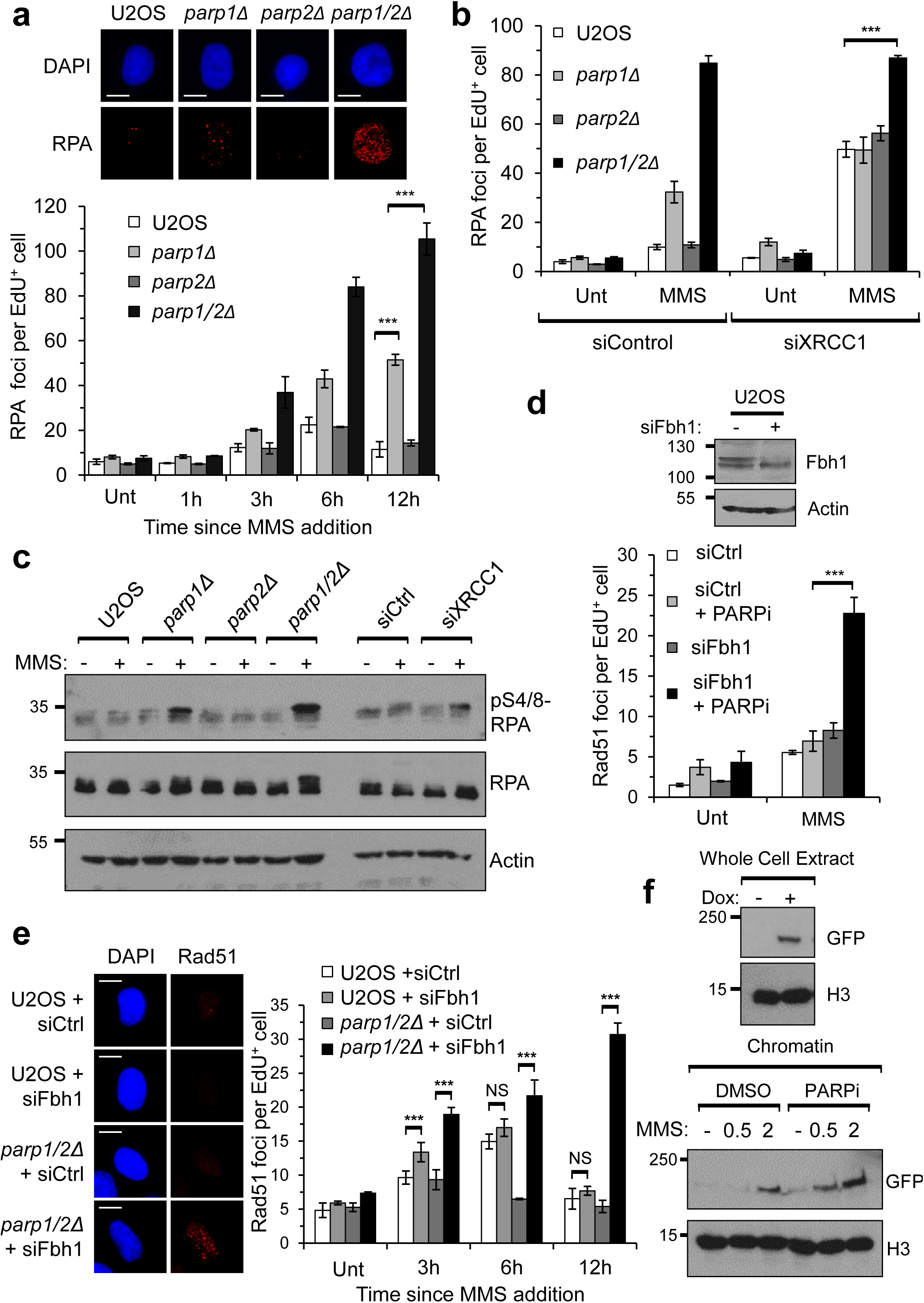
PARP1 and PARP2 are required to stabilise Rad51 at stalled and/or damaged replication forks. **a.** The indicated cell lines were treated with 1 μM EdU (Unt), or with 1 μM EdU and 0.5 mM MMS for 1 hour before recovery in fresh media, and RPA foci quantified in EdU-positive cells. Times indicated show hours since addition of EdU and MMS. Representative images of EdU-positive nuclei 12 hours after addition of MMS are shown (upper panel). Scale bars represent 10 μm. Error bars represent the SEM from three independent experiments. **b.** The indicated cell lines transfected with control or XRCC1 siRNA were treated as in (A) and RPA nuclear foci analysed in EdU positive cells. Cells were fixed 12 hours after addition of EdU and MMS. Error bars represent the SEM from three independent experiments. **c.** The indicated cell lines, with or without transfection of control or XRCC1 siRNA, were left untreated (-) or exposed to 0.5 mM MMS for 1 hour before recovery in fresh media for 12 hours (+). Western blotting of whole cell extracts was performed with the indicated antibodies. **d.** U2OS cells transfected with control or Fbh1 siRNA -/+ olaparib (PARPi), were treated as in (A) and Rad51 nuclear foci analysed in EdU positive cells. Cells were fixed 12 hours after addition of EdU and MMS. Knockdown of Fbh1 was confirmed by Western blotting. Error bars represent the SEM from three independent experiments. **e.** U2OS or *parp1/2∆* cells transfected with control or Fbh1 siRNA were treated as in (A) and Rad51 nuclear foci analysed in EdU positive cells. Times indicated show hours since addition of EdU and MMS. Representative images of EdU-positive nuclei 12 hours after MMS addition are shown. Scale bars represent 10 μm. Error bars represent the SEM from three independent experiments. **f.** Flp-In T-Rex HT1080 cells expressing FE-PALB2, with or without Olaparib (PARPi), were left untreated (-) or exposed to 0.5 mM MMS for 1 hour before recovery in fresh media (+). Extracts were prepared 6 hours after MMS addition. Western blotting of whole cell and chromatin extracts was performed with the indicated antibodies.

Rad51 protects nascent DNA from degradation at stalled replication forks and loss of factors required for loading Rad51 onto DNA, such as BRCA2 and FANCD2 results in extensive nucleolytic degradation of DNA ends^38, 39, 40^. Therefore, one plausible explanation for the extensive ssDNA formation in MMS-treated *parp1/2∆* cells is that loading of factors required to assemble Rad51 onto resected DNA ends is defective. Recently, however, several DNA helicases have been identified that control the stability of a Rad51 nucleofilament^41, 42, 43^. Therefore, an alternative interpretation of our observations is that PARPs regulate Rad51 stability at sites of stalled/damaged replication forks, as opposed to recruitment of Rad51 to sites of replication stress *per se*. To assess these two possibilities, we depleted F-Box DNA helicase 1 (Fbh1), a factor capable of displacing Rad51 from nucleofilaments and suppressing HR^43, 44^. Depletion of Fbh1 results in a slight increase in Rad51 foci formation in replicating U2OS cells at an early time point following exposure to MMS, although they decay to a similar extent to that observed in control cells (Fig. 5e). Strikingly, however, it restores Rad51 foci formation in PARPi treated U2OS, RPE-1 and MRC5 cells (Fig. 5d and Supplementary Fig. 11), or *parp1/2∆* U2OS cells (Fig. 5e). Additionally, PARPi do not significantly decrease the levels of PALB2 enriched in chromatin following exposure of cells to MMS, indicating that the ability of this accessory factor required for loading Rad51 at sites of DNA damage is not affected by PARP status (Fig. 5f). We observe that loss of PARP1 and PARP2 also compromises Rad51 foci formation in response to hydroxyurea (HU), and that this defective Rad51 foci formation can be similarly restored by depletion of Fbh1 (Supplementary Fig. 12), indicating that this observation is not restricted to MMS-induced replication stress. Furthermore, and consistent with problematic Rad51 retention, replication fork stability is also severely compromised after prolonged HU exposure upon combined loss of PARP1 and PARP2 (Supplementary Fig. 12). Taken together, these data support the notion that PARP1 and PARP2 do not regulate the recruitment of Rad51 to stalled and/or damaged replication forks, but instead act to stabilise Rad51 nucleofilaments to promote repair of replication associated DNA damage induced by MMS.

## DISCUSSION

Our observations that *parp1∆* cells can be further sensitised to MMS by PARPi indicate an additional PARP(s) is required for the cellular response to DNA base alkylation. Our data indicate PARP2 performs this role and we uncover strong redundancy between PARP1 and PARP2 in the cellular response to DNA base alkylation. PARP2 was originally proposed as a possible ARTD responsible for residual DNA damage-induced ADP-ribosylation^45, 46^. Consistent with this hypothesis, we observe that residual MMS-induced ADP-ribosylation in *parp1∆* cells is significantly reduced by disruption of *parp2* (Fig. 1d). Moreover, disruption of *parp2* in a *parp1∆* background further sensitises cells to MMS and *parp1/2∆* cells exhibit a delay in resolution of this damage compared to U2OS, *parp1∆* and *parp2∆* cells (Fig. 1e,f). Given PARP2 is recruited to laser induced DNA damage sites after PARP1^47^, it is tempting to speculate that PARP2 lies downstream of PARP1 in the SSBR pathway. However, it is noteworthy that disruption of *parp2* does not further sensitise *parp1∆* cells to SSBs induced by H_2_O_2_ (Supplementary Fig. 2b) or delay the resolution of DNA breaks induced by this agent, even in the absence of PARP1^48, 49^. Moreover, effective resolution of breaks generated during BER is apparent in the absence of PARP1 or PARP2, arguing against an absolute requirement for either of these enzymes alone in this process. Together, these data support redundant roles for PARP1 and PARP2 in BER, as opposed to SSBR more generally.

It is well documented that PARPi are toxic to cells with defects in HR^24^. However, given currently available PARPi target multiple ARTDs^22, 23^, which PARP is the most effective to target in synthetic lethal strategies with HR-deficiency remains an open question. Our findings indicate that inhibition of HR is toxic to cells disrupted in *PARP1* (Fig. 2b). Cells are able to tolerate HR inhibition equally well in the presence or absence of PARP3 (Fig. 2c), in keeping with the observations that AZD2461, a compound that inhibits PARP1 and PARP2 but not PARP3, effectively causes *BRCA*-mutated tumour regression^50^. PARP2 status does not dramatically affect the ability of cells to survive HR inhibition, even in the absence of PARP1 (Fig. 2b). This is surprising given the redundancy between PARP1 and PARP2 in repair of DNA base alkylation induced by MMS (Fig. 1). However, it should be noted that we do not observe redundancy between PARP1 and PARP2 in sensitivity of cells to H_2_O_2_, an agent that induces oxidative DNA damage that might more accurately reflect the type of DNA lesion encountered by replication forks in a physiological setting. We also observe that depletion of the key SSBR factor XRCC1 has little impact on the ability of cells to tolerate HR inhibition (Fig. 2d), supporting the model that PARPi toxicity is driven by trapping the PARPs at SSBs to elicit replication blockage, as opposed to SSBR defects *per se*^25, 51^. However, *parp1∆* cells are clearly sensitive to B02, indicating PARP trapping is not the only factor that drives toxicity with HR inhibition. Importantly, our data clearly indicate that disruption of PARP1 is more toxic with HR inhibition than loss of either PARP2 or PARP3. Given PARP2 is essential for hematopoietic stem/progenitor cell survival^52^, our observations would support the development of PARP1 selective inhibitors that may limit off-target effects associated with broader specificity compounds.

Taken together, our data support an additional role of PARP1 and PARP2 in allowing cells to tolerate DNA base alkylation beyond their regulation of BER/SSBR. We clearly identify that whilst PARP1 and PARP2 are required for canonical BER, they are also required to resolve replication associated DNA damage during DNA synthesis. Our data identifying that decay of γ-H2AX foci is delayed in *parp1∆* cells indicate that this is attributable, in part, to PARP1. Strikingly however, this phenotype is exacerbated in *parp1/2∆* cells, indicating further redundancy between PARP1 and PARP2 in repair of replication associated DNA damage induced by MMS. S-phase associated γ-H2AX foci induced by MMS are suppressed by depletion of *N*-methylpurine DNA glycosylase (MPG) or AP-endonuclease, the enzymes that remove the methylated base and perform the strand incision step to initiate BER^30^. This supports the notion that PARP1 and PARP2 are required to repair S-phase associated DNA DSBs that arise as a consequence of replication forks encountering SSB repair intermediates generated during BER. Depletion of XRCC1 does not result in a similar persistence of MMS-induced γ-H2AX foci (Fig. 3d,e), indicating that XRCC1 does not contribute significantly to replication associated repair in this context. Importantly, however, this also supports a model whereby the inability to resolve DNA damage in the absence of PARP1 and PARP2 is independent of BER, identifying a direct role of these enzymes in repair of MMS-induced DNA damage during S-phase. Thus, PARP1 and PARP2 are critical for two separate but linked aspects of repair of alkylated DNA base damage; one through regulating BER, in addition to repair of replication associated DNA strand breaks generated as a consequence of unresolved SSB intermediates resulting in replication stress.

HR protects cells from replication associated DSBs induced by MMS^53^ and PARP1 has been implicated in the slowing and/or restart of replication forks in response to genotoxins^11, 12, 13, 54^. However, whether PARP1 regulates HR at stalled/damaged replication forks, how this is achieved, and the role of PARP2 in these events is unclear. Our data demonstrate a critical requirement for PARP1 and PARP2 in stabilising the Rad51 nucleofilament at damaged replication forks. Whilst an induction of Rad51 foci is evident in replicating U2OS, *parp1∆* and *parp2∆* cells exposed to MMS, their decay is delayed in the absence of PARP1 (Fig. 4b), indicating either a problem in resolving HR intermediates, or increased levels of DNA damage being channelled through HR. Importantly, we observe that XRCC1 depletion results in persistent Rad51 foci in several different cell lines (Fig. 4b and Supplementary Fig. 9), supporting the notion that defective SSBR/BER results in elevated levels of DNA damage that are channelled through Rad51-dependent repair mechanisms. Strikingly, however, we observe that in the absence of PARP1 and PARP2, replication fork stalling and assembly of Rad51 at sites of replication stress induced by MMS are compromised (Fig. 4a,b and Supplementary Fig. 10). Given the critical role of Rad51 in the stabilisation, repair and restart of damaged replication forks, this provides an explanation for the persistent levels of DNA damage in these cells.

The ability to form RPA foci in response to MMS independent of PARP status would argue against DNA end-resection being point at which PARP1 and PARP2 regulate assembly of Rad51 into nucleofilaments (Fig. 5a,b). Indeed, we observe significantly elevated levels of RPA foci and pS4/8 RPA in *parp1/2∆* cells, despite their inability to form Rad51 foci in response to MMS (Fig. 5a-c). We also observe a decrease in Rad51 foci in *parp1/2∆* cells exposed to HU that is accompanied by an increase in replication fork degradation (Supplementary Fig. 12), indicating the link between PARP1 and PARP2, Rad51 stabilisation in chromatin and replication fork stability may be applicable more generally. These observations are consistent with the reported role of Rad51 in protecting stalled and damaged replication forks from Mre11 and DNA2-dependent nucleolytic attack^39, 40, 55, 56^ and can be interpreted either as a defect in recruitment of Rad51 to impaired replication forks and/or stabilisation of the Rad51 nucleofilament. Importantly, depletion of Fbh1, an anti-recombination helicase that destabilises Rad51 nucleofilaments^43, 44, 57^, can restore MMS and HU-induced Rad51 foci in *parp1/2∆* cells or U2OS cells treated with olaparib (Fig. 5d,e). Whilst this observation could reflect a shift in the equilibrium between Rad51 loading and removal from nucleofilaments, depletion of Fbh1 cannot restore Rad51 foci in BRCA2 defective cells, arguing against a role for PARP1/PARP2 in BRCA2-dependent loading of Rad51 at damaged replication forks^55^. Consistent with this hypothesis, enrichment of PALB2 into chromatin following exposure of cells to MMS is not affected by olaparib (Fig. 5f), a PARPi that targets both PARP1 and PARP2 and can suppress formation of Rad51 foci in response to MMS (Fig. 4c). Instead, our data argue that PARPs act downstream of BRCA2/PALB2 and function to stabilise Rad51 at the sites of stalled/damaged replication forks.

In summary, our data point to the following model of how PARP1 and PARP2 regulate repair of alkylated DNA damage by two independent, but intrinsically linked mechanisms (Figure 6): PARP1 and PARP2 function redundantly in repair of alkylated DNA base damage by promoting BER. However, removal of the base and subsequent cleavage by APE1 results in a subset of SSBs being converted to DSBs when they are encountered by active replication forks. Under normal circumstances, limited nucleolytic resection allows assembly of Rad51 at damaged replication forks. PARP1 and PARP2 regulate this process by opposing the role of Fbh1, either directly or indirectly, in destabilising Rad51 at damaged replication forks. Therefore, under normal circumstances PARP1 and PARP2 function redundantly to stabilise Rad51 nucleofilaments, facilitating HR-dependent replication fork repair. However, loss of PARP1 and PARP2 leads to defective BER, resulting in elevated levels of SSBs and consequently replication associated DNA damage. It also relieves the opposing role of PARPs on Fbh1, resulting in disassembly of the Rad51 nucleofilament, uncontrolled resection at stalled and/or damaged replication forks, and an inability to repair DNA damage.

**Figure 6:**
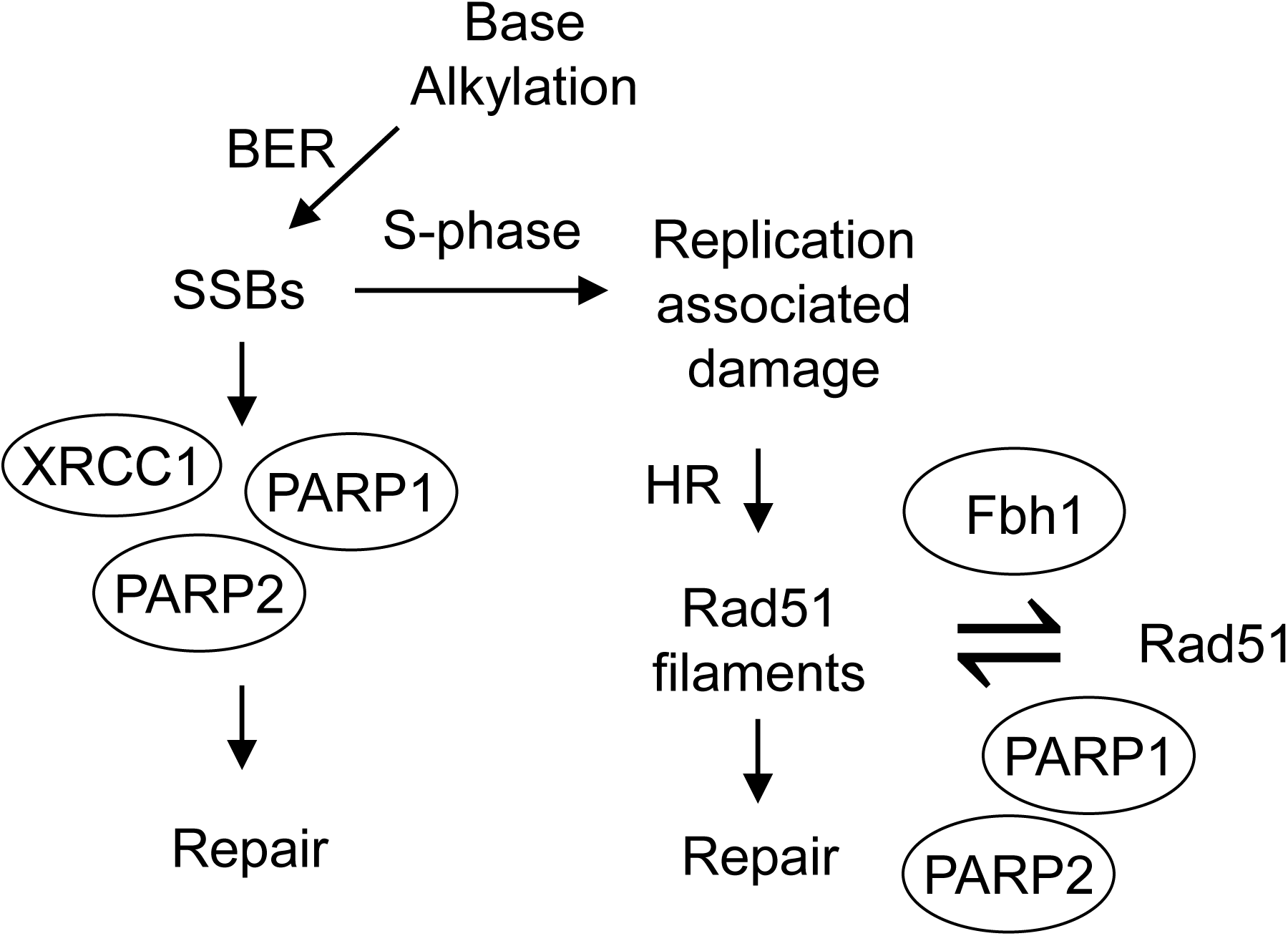
PARP1 and PARP2 contribute to the resolution of DNA damage caused by base alkylation through two mechanisms. Alkylated bases are processed into DNA single strand breaks during BER and their repair is accelerated by XRCC1, PARP1 and PARP2. During S-phase, active replication forks can collide with unrepaired SSBs, or SSB repair intermediates, generating replication-associated DNA damage. This damage can be resolved through homologous recombination, which requires the formation of Rad51 filaments. PARP1 and PARP2 also contribute to this mechanism by antagonising the anti-recombinogenic activity of Fbh1, thus stabilising Rad51 nucleofilaments.

## METHODS

### Cell culture and siRNA transfections

U2OS cells were cultured in DMEM supplemented with 10% FBS and 1% Pen/Strep. The identity of U2OS cells and their derivatives were confirmed by STR profiling. Flp-In T-Rex HT1080 cells expressing FE-PALB2 were a kind gift of F. Esashi (Dunn School of Pathology, University of Oxford), and were cultured in DMEM as above, supplemented with 100 μg/mL Hygromycin and 10 μg/mL Blasticidin. All cell lines were tested for the absence of mycoplasma contamination.

Cells were transfected with 50 nM siRNA using Dharmafect-1 (Dharmacon), transfected again after 24 hours, then allowed to recover for 48 hours before use. siRNA SMARTpools were obtained from Dharmacon, either ON-Target Plus (PARP2, PARP3 and Fbh1) or siGenome (XRCC1), and the appropriate non-targetting control SMARTpool used for negative controls.

### CRISPR cell line generation

Pairs of CRSIPR gRNAs were designed using the MIT CRISPR design tool (http://crispr.mit.edu/), and cloned into pX462 or pX462 v2 (pSpCas9n(BB)-2A-Puro).

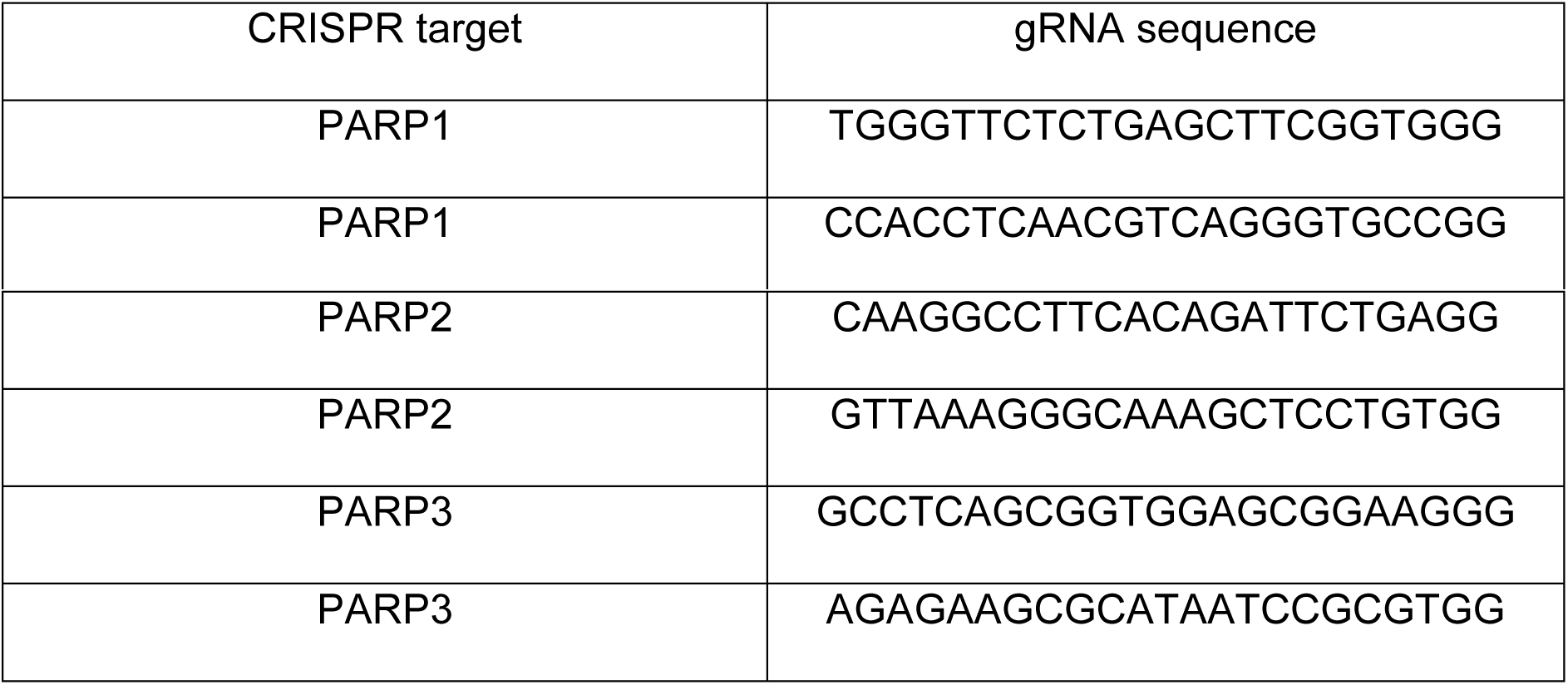

U2OS cells were transfected with the appropriate gRNA constructs using Turbofect according to manufacturer’s instructions, selected with 2-4 μg/mL puromycin for 24 hours, then plated at low density. Clonal colonies were isolated and expanded, then screened for lack of the relevant protein by Western blot. Gene disruption was confirmed using PCR and sequencing across the CRISPR target site to characterise the mutations generated. Genomic DNA was prepared from candidate cell lines by resuspending cells in DNA Extraction Buffer (10 mM Tris-HCL pH 8.0, 1 mM EDTA, 25 mM NaCl and 200 μg/mL Proteinase K), heating to 65 °C for 30 minutes, then 95 °C for 2 minutes. After PCR amplification, bands were cloned into pJet1.2 (ThermoFisher) before analysis by Sanger sequencing. The *parp1/2∆* cells were generated by disrupting PARP1 in a *parp2∆* cell line.

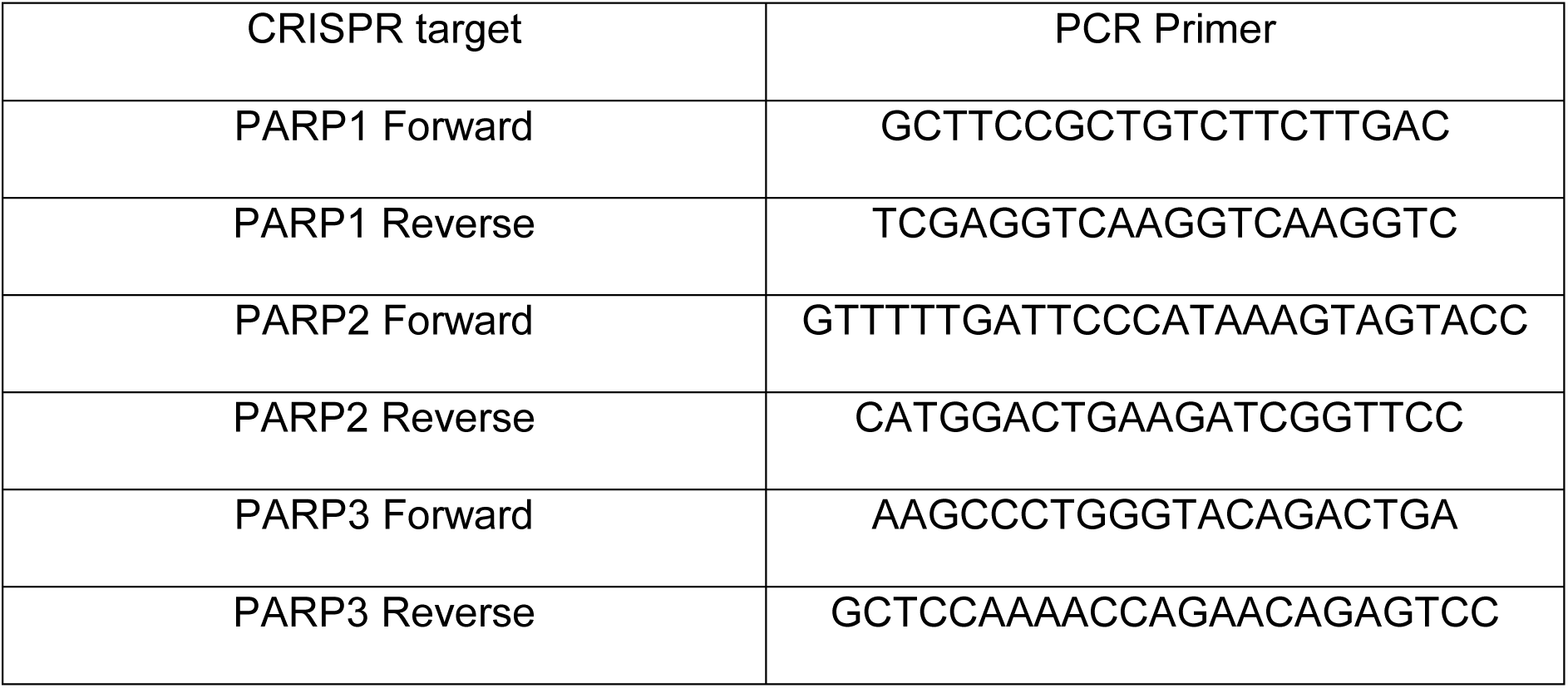

### Antibodies and Chemicals

MMS (Sigma-Aldrich) or Hydrogen Peroxide (Sigma-Aldrich) were freshly diluted into cell culture media directly before use. Cells were treated with the indicated concentrations for one hour, washed extensively with PBS, and allowed to recover in fresh media as necessary. Olaparib (Cambridge Bioscience) was used at 10 μM, and cells were treated with Olaparib 1 hour prior to DNA damage. B02 (Sigma-Aldrich) was used at the indicated concentrations. EdU was used at a final concentration of 1 μM, and added to cells simultaneously with DNA damaging agents. Nu7441 (Tocris Bioscience) was used at a final concentration of 5 μM. CldU (Sigma-Aldrich) and IdU (Sigma-Aldrich) were used at final concentrations of 25 μM or 250 μM respectively under conditions as described for the fibre experiment. To induce FE-PALB2 expression, Flp-In T-Rex HT1080 cells containing FE-PALB2 were treated with 5 μg/mL Doxycyclin for 24 hours.

Antibodies used in this study were anti-PARP1 (Bio-Rad, MCA1522G), anti-PARP2 (Enzo, ALX-804-639L), anti-PARP3 (A kind gift of F. Dantzer; Institut de Recherche de l'Ecole de Biotechnologie de Strasbourg, France), anti-Poly-ADP-Ribose binding reagent (Millipore, MABE1031), anti-XRCC1 (Abcam, ab1838), anti-Actin (Santa Cruz, SC-1615), anti-γH2AX (Millipore, JBW301), anti-Rad51 (7946; a kind gift of F. Esashi; Dunn School of Pathology, University of Oxford), anti-RPA32 (Bethyl, A300-244A), anti-RPA32 S4/8 (Bethyl, A300-245A), anti-GFP (Roche, 11814460001), anti-Fbh1 (Abcam, ab58881), mouse anti-BrdU (clone B44 recognising IdU, Becton Dickinson, 347580) and rat anti-BrdU (clone BU1/75 recognising CldU, Bio-Rad, OBT0030G).

### Western Blotting and Cellular Fractionation

Whole cell extracts were prepared by boiling cells in 1x SDS loading buffer for 5 minutes.

Chromatin extracts were prepared by washing cells in PBS before resuspending in extraction buffer 1 (10 mM Tris-HCl pH 7.5, 150 mM NaCl, 1.5 mM MgCl_2_, 340 mM sucrose, 10 % glycerol, 1 mM DTT, 1 x Sigma protease inhibitors and 0.1% Triton X-100). The samples were incubated on ice for 10 minutes, then centrifuged at 18,000g for 5 minutes, then the pellet resuspended in extraction buffer 2 (3 mM EDTA, 0.2 mM EGTA, 1 mM DTT, 1 x Sigma Protease Inhibitors) and incubated on ice for 30 minutes. Samples were centrifuged again at 1800g for 5 minutes, and the pellet resuspended in 1x SDS loading buffer and boiled.

### Clonogenic survival assays

U2OS cells were plated into 6-well plates and left overnight. The following day, cells were exposed to DNA damaging agents for 1 hour, washed extensively with PBS then allowed to recover in fresh media for 10-14 days. For synthetic lethality experiments using Rad51 inhibition, cells were exposed to B02 and/or Olaparib or Nu7441 for 48 hours at the indicated doses, before washing and recovery in fresh media. Cells were then fixed in 100% methanol for 20 minutes at -20 °C and stained with 0.5% Crystal Violet (Sigma Aldrich) for 20 minutes. Colonies of more than 50 cells were scored as viable.

### Immunofluorescence

Cells were plated onto glass coverslips and allowed to attach overnight. Following DNA damage treatment, cells were fixed with 4% paraformalaldehyde for 20 minutes at 4 °C. Where necessary, prior to fixation, soluble protein was pre-extracted using 0.5% Triton X-100 in PBS for 5 minutes at 4 °C. Cells were then permeablisied in 0.5% Triton X-100 in PBS for 5 minutes, and blocked in 3% BSA for 30 minutes. EdU was stained by incubation in reaction buffer (100 mM Tris-Cl pH 8.0, 4 mM CuSO4, 50 mM sodium ascorbate, 20 μM azide dye) for 30 minutes. Coverslips were stained with primary antibody (2 hours, room temperature), washed extensively in PBS-T, and stained with fluorescently labelled secondary antibody (1 hour, room temperature). Following further PBS-T washing, cells were mounted onto slides in Vectashield containing DAPI (Vector Laboratories). Samples were visualised using a Zeiss IX70 and 100x oil immersion objective lens. Each experiment was independently repeated at least three times, with a total of at least 100 cells counted per condition. Images were processed in ImageJ.

### DNA fibre analysis

Replication fork degradation assays following exposure of cells to HU were performed as previously described^55^. To assess replication fork stalling, cells were pulse-labelled with 25 μM CldU for 10 min prior to 40 min co-incubation with 0.5 mM MMS. MMS-treated cells were washed trice with PBS and released into fresh media containing 250 μM IdU for 0.5, 1 and 2 h before harvesting. DNA fibre spreads were prepared by spotting 2 μL of cell solution (5 × 10^5^ cells per mL PBS) onto microscope slides followed by lysis with 7 μL of lysis buffer (0.5 % SDS, 200 mM Tris-HCl pH 7.4 and 50 mM EDTA). For spreading, slides were tilted and subsequently fixed in methanol/acetic acid (3:1). HCl-treated fibre spreads were treated with rat anti-BrdU (binding to CldU, clone BU1/75, Bio-Rad, 1:500) and mouse anti-BrdU (binding to IdU, clone B44, Becton Dickinson, 1:500) for 1 h and subsequently fixed with 4 % paraformaldehyde for 10 min to increase staining intensity. Afterwards, slides were incubated with anti-rat IgG AlexaFluor 555 and anti-mouse IgG AlexaFluor 488 (Molecular Probes) for 1.5 h. Images were acquired on a Nikon E600 microscope using a Nikon Plan Apo x 60 (1.3 numerical aperture) oil lens, a Hamamatsu digital camera (C4742-95) and the Volocity acquisition software (Perkin Elmer). Replication structures were analysed using ImageJ (http://rsbweb.nih.gov/ij/) applying the cell counting plugin. At least 300 replication structures were assessed per condition.

### Alkaline comet assays

The method was modified from a previously described procedure to detect DNA strand breaks and alkali labile sites^58, 59^. Cells were diluted to a concentration of 1×10^5^ / ml in PBS. The cell suspension was mixed with an equal volume of warm 2% agarose solution (Type VII). 500 μl of the suspension was spread evenly onto polylysine coated slides that are pre-coated with 0.3% agarose (Type 1). For cell lysis, the slides were soaked in 0.3 M NaOH/1 M NaCl/0.1 % N-lauroylsarcosine, pH 11.5 in the dark for 1 h followed by two 30 min washes in 0.3 M NaOH, 2 mM EDTA. Slides were electrophoresed at 75 mA for 30 min in the same buffer at pH 11.5. Following electrophoresis slides were flooded with neutralisation buffer (Tris-Cl pH7.5) for 30 minutes. Slides were dried overnight in the dark, and subsequently rehydrated in distilled water for 30 min in the dark. Slides were stained with SYBR gold (1:10,000 in water) for 20 min and then rinsed in water before leaving to dry overnight. The percentage DNA in the tail and Olive moment were determined using OpenComet. The area and mean pixel intensity of the head and the tail of the comets were measured to determine the percentage DNA in the tail for the individual cell. For each experimental sample 300 cells per time point (from duplicate slides) were analysed and the mean Olive Moment was determined.

### Quantification and statistical analysis

Statistical significance was determined on at least 3 biological replicates using a two-tailed Student’s t-test and is indicated as: NS, not significant; * p < 0.05; ** p<0.01; *** p<0.001.

### Data Availability

All data generated or analysed during this study are included in this published article (and its supplementary information files).

## ACKNOWLEDGEMENTS

NDL’s laboratory is supported by the Medical Research Council (www.mrc.ac.uk; MR/L000164/1 and MR/P018963/1). GR was supported by a Wellcome Trust Studentship (102348/Z/13/Z). AO is funded by the Oxford Experimental Cancer Medicine Centre and ALP by a German Research Foundation Fellowship (Pl 1300/1-1). GSS is supported by a Cancer Research UK programme grant (C17183/A23303) and MRH by an MRC Career Development Fellowship (MR/P009085/1) and University of Birmingham Fellowship.

## AUTHOR CONTRIBUTIONS

All experiments were performed by GR with the exception of comets assays (AO and PJM), DNA Fibre analysis (ALP and EP) and fork protection assays (MRH and GSS). GR and NDL, PJM and ALP wrote the manuscript.

## MATERIALS AND CORRESPONDANCE

Correspondence and materials requests should be made to Nick Lakin.

## REFERENCES

1. Jackson SP, Bartek J. The DNA-damage response in human biology and disease. Nature 461, 1071-1078 (2009).

2. Polo SE, Jackson SP. Dynamics of DNA damage response proteins at DNA breaks: a focus on protein modifications. Genes Dev 25, 409-433 (2011).

3. Gibson BA, Kraus WL. New insights into the molecular and cellular functions of poly(ADP-ribose) and PARPs. Nat Rev Mol Cell Biol 13, 411-424 (2012).

4. Hottiger MO, Hassa PO, Luscher B, Schuler H, Koch-Nolte F. Toward a unified nomenclature for mammalian ADP-ribosyltransferases. Trends Biochem Sci 35, 208-219 (2010).

5. Hottiger MO. Nuclear ADP-Ribosylation and Its Role in Chromatin Plasticity, Cell Differentiation, and Epigenetics. Annu Rev Biochem 84, 227-263 (2015).

6. Caldecott KW. Single-strand break repair and genetic disease. Nat Rev Genet 9, 619-631 (2008).

7. Caldecott KW, Aoufouchi S, Johnson P, Shall S. XRCC1 polypeptide interacts with DNA polymerase beta and possibly poly(ADP-ribose) polymerase, and DNA ligase III is a novel molecular 'nick-sensor' *in vitro*. Nucleic Acids Res 24, 4387-4394 (1996).

8. El-Khamisy SF, Masutani M, Suzuki H, Caldecott KW. A requirement for PARP-1 for the assembly or stability of XRCC1 nuclear foci at sites of oxidative DNA damage. Nucleic Acids Res 31, 5526-5533 (2003).

9. Masson M, Niedergang C, Schreiber V, Muller S, Menissier de Murcia J, deMurcia G. XRCC1 is specifically associated with poly(ADP-ribose) polymerase and negatively regulates its activity following DNA damage. Mol Cell Biol 18, 3563-3571 (1998).

10. Okano S, Lan L, Caldecott KW, Mori T, Yasui A. Spatial and temporal cellular responses to single-strand breaks in human cells. Mol Cell Biol 23, 3974-3981 (2003).

11. Bryant HE, et al. PARP is activated at stalled forks to mediate Mre11-dependent replication restart and recombination. Embo J 28, 2601-2615 (2009).

12. Sugimura K, Takebayashi S, Taguchi H, Takeda S, Okumura K. PARP-1 ensures regulation of replication fork progression by homologous recombination on damaged DNA. J Cell Biol 183, 1203-1212 (2008).

13. Yang YG, Cortes U, Patnaik S, Jasin M, Wang ZQ. Ablation of PARP-1 does not interfere with the repair of DNA double-strand breaks, but compromises the reactivation of stalled replication forks. Oncogene 23, 3872-3882 (2004).

14. Audebert M, Salles B, Calsou P. Involvement of poly(ADP-ribose) polymerase-1 and XRCC1/DNA ligase III in an alternative route for DNA double-strand breaks rejoining. J Biol Chem 279, 55117-55126 (2004).

15. Wang M, Wu W, Rosidi B, Zhang L, Wang H, Iliakis G. PARP-1 and Ku compete for repair of DNA double strand breaks by distinct NHEJ pathways. Nucleic Acids Res 34, 6170-6182 (2006).

16. Luijsterburg MS, et al. PARP1 Links CHD2-Mediated Chromatin Expansion and H3.3 Deposition to DNA Repair by Non-homologous End-Joining. Mol Cell 61, 547-562 (2016).

17. Boehler C, et al. Poly(ADP-ribose) polymerase 3 (PARP3), a newcomer in cellular response to DNA damage and mitotic progression. Proc Natl Acad Sci U S A 108, 2783-2788 (2011).

18. Couto CA, et al. PARP regulates nonhomologous end joining through retention of Ku at double-strand breaks. J Cell Biol 194, 367-375 (2011).

19. Rulten SL, et al. PARP-3 and APLF function together to accelerate nonhomologous end-joining. Mol Cell 41, 33-45 (2011).

20. Menissier de Murcia J, et al. Functional interaction between PARP-1 and PARP-2 in chromosome stability and embryonic development in mouse. Embo J 22, 2255-2263 (2003).

21. Schreiber V, et al. Poly(ADP-ribose) polymerase-2 (PARP-2) is required for efficient base excision DNA repair in association with PARP-1 and XRCC1. J Biol Chem 277, 23028-23036 (2002).

22. Wahlberg E, et al. Family-wide chemical profiling and structural analysis of PARP and tankyrase inhibitors. Nat Biotechnol 30, 283-288 (2012).

23. Thorsell AG, et al. Structural Basis for Potency and Promiscuity in Poly(ADP-ribose) Polymerase (PARP) and Tankyrase Inhibitors. J Med Chem 60, 1262-1271 (2017).

24. O'Connor MJ. Targeting the DNA Damage Response in Cancer. Mol Cell 60, 547-560 (2015).

25. Pommier Y, O'Connor MJ, de Bono J. Laying a trap to kill cancer cells: PARP inhibitors and their mechanisms of action. Sci Transl Med 8, 362ps317 (2016).

26. Huang F, Mazina OM, Zentner IJ, Cocklin S, Mazin AV. Inhibition of homologous recombination in human cells by targeting RAD51 recombinase. J Med Chem 55, 3011-3020 (2012).

27. Patel AG, Sarkaria JN, Kaufmann SH. Nonhomologous end joining drives poly(ADP-ribose) polymerase (PARP) inhibitor lethality in homologous recombination-deficient cells. Proc Natl Acad Sci U S A 108, 3406-3411 (2011).

28. Dianov GL, Hubscher U. Mammalian base excision repair: the forgotten archangel. Nucleic Acids Res 41, 3483-3490 (2013).

29. Kaina B. Mechanisms and consequences of methylating agent-induced SCEs and chromosomal aberrations: a long road traveled and still a far way to go. Cytogenet Genome Res 104, 77-86 (2004).

30. Ensminger M, Iloff L, Ebel C, Nikolova T, Kaina B, Lobrich M. DNA breaks and chromosomal aberrations arise when replication meets base excision repair. J Cell Biol 206, 29-43 (2014).

31. Brem R, Hall J. XRCC1 is required for DNA single-strand break repair in human cells. Nucleic Acids Res 33, 2512-2520 (2005).

32. Kubota Y, Horiuchi S. Independent roles of XRCC1’s two BRCT motifs in recovery from methylation damage. DNA Repair (Amst) 2, 407-415 (2003).

33. Taylor RM, Thistlethwaite A, Caldecott KW. Central role for the XRCC1 BRCT I domain in mammalian DNA single-strand break repair. Mol Cell Biol 22, 2556-2563 (2002).

34. Horton JK, Watson M, Stefanick DF, Shaughnessy DT, Taylor JA, Wilson SH. XRCC1 and DNA polymerase beta in cellular protection against cytotoxic DNA single-strand breaks. Cell research 18, 48-63 (2008).

35. Zellweger R, et al. Rad51-mediated replication fork reversal is a global response to genotoxic treatments in human cells. J Cell Biol 208, 563-579 (2015).

36. Henry-Mowatt J, et al. XRCC3 and Rad51 modulate replication fork progression on damaged vertebrate chromosomes. Mol Cell 11, 1109-1117 (2003).

37. Budzowska M, Kanaar R. Mechanisms of dealing with DNA damage-induced replication problems. Cell biochemistry and biophysics 53, 17-31 (2009).

38. Hashimoto Y, Ray Chaudhuri A, Lopes M, Costanzo V. Rad51 protects nascent DNA from Mre11-dependent degradation and promotes continuous DNA synthesis. Nat Struct Mol Biol 17, 1305-1311 (2010).

39. Schlacher K, Christ N, Siaud N, Egashira A, Wu H, Jasin M. Double-strand break repair-independent role for BRCA2 in blocking stalled replication fork degradation by MRE11. Cell 145, 529-542 (2011).

40. Schlacher K, Wu H, Jasin M. A distinct replication fork protection pathway connects Fanconi anemia tumor suppressors to RAD51-BRCA1/2. Cancer Cell 22, 106-116 (2012).

41. Bugreev DV, Yu X, Egelman EH, Mazin AV. Novel pro- and anti-recombination activities of the Bloom’s syndrome helicase. Genes Dev 21, 3085-3094 (2007).

42. Hu Y, et al. RECQL5/Recql5 helicase regulates homologous recombination and suppresses tumor formation via disruption of Rad51 presynaptic filaments. Genes Dev 21, 3073-3084 (2007).

43. Chu WK, Payne MJ, Beli P, Hanada K, Choudhary C, Hickson ID. FBH1 influences DNA replication fork stability and homologous recombination through ubiquitylation of RAD51. Nat Commun 6, 5931 (2015).

44. Fugger K, et al. Human Fbh1 helicase contributes to genome maintenance via pro- and anti-recombinase activities. J Cell Biol 186, 655-663 (2009).

45. Ame JC, et al. PARP-2, a novel mammalian DNA damage-dependent poly(ADP-ribose) polymerase. J Biol Chem 274, 17860-17868 (1999).

46. Shieh WM, et al. Poly(ADP ribose) polymerase null mouse cells synthesize ADP ribose polymers. J Biol Chem 273, 30069-30072 (1998).

47. Mortusewicz O, Ame JC, Schreiber V, Leonhardt H. Feedback-regulated poly(ADP-ribosyl)ation by PARP-1 is required for rapid response to DNA damage in living cells. Nucleic Acids Res 35, 7665-7675 (2007).

48. Fisher AE, Hochegger H, Takeda S, Caldecott KW. Poly(ADP-ribose) polymerase 1 accelerates single-strand break repair in concert with poly(ADP-ribose) glycohydrolase. Mol Cell Biol 27, 5597-5605 (2007).

49. Hanzlikova H, Gittens W, Krejcikova K, Zeng Z, Caldecott KW. Overlapping roles for PARP1 and PARP2 in the recruitment of endogenous XRCC1 and PNKP into oxidized chromatin. Nucleic Acids Res, (2016).

50. Oplustil O'Connor L, et al. The PARP Inhibitor AZD2461 Provides Insights into the Role of PARP3 Inhibition for Both Synthetic Lethality and Tolerability with Chemotherapy in Preclinical Models. Cancer Res 76, 6084-6094 (2016).

51. Murai J, et al. Trapping of PARP1 and PARP2 by Clinical PARP Inhibitors. Cancer Res 72, 5588-5599 (2012).

52. Farres J, et al. Parp-2 is required to maintain hematopoiesis following sublethal gamma-irradiation in mice. Blood 122, 44-54 (2013).

53. Nikolova T, Ensminger M, Lobrich M, Kaina B. Homologous recombination protects mammalian cells from replication-associated DNA double-strand breaks arising in response to methyl methanesulfonate. DNA Repair (Amst) 9, 1050-1063 (2010).

54. Berti M, et al. Human RECQ1 promotes restart of replication forks reversed by DNA topoisomerase I inhibition. Nat Struct Mol Biol 20, 347-354 (2013).

55. Higgs MR, et al. BOD1L Is Required to Suppress Deleterious Resection of Stressed Replication Forks. Mol Cell 59, 462-477 (2015).

56. Karanja KK, Lee EH, Hendrickson EA, Campbell JL. Preventing over-resection by DNA2 helicase/nuclease suppresses repair defects in Fanconi anemia cells. Cell Cycle 13, 1540-1550 (2014).

57. Simandlova J, et al. FBH1 helicase disrupts RAD51 filaments in vitro and modulates homologous recombination in mammalian cells. J Biol Chem 288, 34168-34180 (2013).

58. Olive PL, Banath JP, Durand RE. Heterogeneity in radiation-induced DNA damage and repair in tumor and normal cells measured using the “comet” assay. Radiat Res 122, 86-94 (1990).

59. Spanswick VJ, Hartley JM, Hartley JA. Measurement of DNA interstrand crosslinking in individual cells using the Single Cell Gel Electrophoresis (Comet) assay. Methods Mol Biol 613, 267-282 (2010).

